# Composite Certainty: Addressing Metric Degeneracy in Parameter Inference for Model-Based Diagnostics

**DOI:** 10.64898/2026.05.09.724027

**Authors:** Amruta Koshe, Ehsan Sobhani Tehrani, Kian Jalaleddini, Hamid Motallebzadeh

## Abstract

Quantifying the diagnostic dispersion of inferred parameter distributions is a challenge in uncertainty-aware modeling. Scalar summaries such as credible interval width are topology-blind; fundamentally different posterior morphologies can yield identical scores, obscuring whether a parameter is precisely estimated or constrained to a range. We propose a Composite Certainty Framework that addresses this metric degeneracy by aggregating five complementary uncertainty metrics including interquartile range, standard deviation, full width at half maximum, Shannon entropy, and mass width. These metrics are aggregated through non-parametric Borda rank voting into a single, unitless consensus certainty score. Applied to a simulation-based inference pipeline for a finite-element model of the human middle ear tuned to cadaveric acoustic measurements, the framework reveals parameter-specific identifiability profiles invisible to any individual metric. It produces two actionable clinical thresholds: (1) the maximum tolerable measurement noise for reliable parameter recovery, and (2) the minimum simulation budget for posterior convergence. We demonstrated that no single metric captures all aspects of posterior dispersion, as spread-based metrics and entropy diverge systematically for clinically critical parameters, whereas their aggregation produces a consensus reflecting genuine diagnostic certainty. The framework is generalizable to any model-based diagnostic pipeline where posterior distribution not merely its coverage, but determines clinical certainty.

## I. INTRODUCTION

Computational biomechanics is undergoing a fundamental paradigm shift from deterministic models toward probabilistic frameworks that acknowledge the inherent uncertainty in biological systems (Cranmer et al., 2020; Viceconti et al., 2020). While traditional models seek a single “best-fit” parameter set, modern machine learning (ML) approaches, such as simulation-based inference (SBI), provide full posterior probability distribution functions (PDF) representing the range of all plausible physical states (Motallebzadeh et al., 2025). These posterior PDFs encode two fundamentally distinct sources of uncertainty: (1) epistemic uncertainty, arising from limited training data and/or measurement noise, which are reducible in principle; and (2) aleatoric uncertainty, arising from intrinsic non-identifiability in the forward model and irreducible regardless of data quality (Der Kiureghian & Ditlevsen, 2009; Kendall & Gal, 2017). These models are increasingly used to drive model-based medical diagnostics, yet a critical gap remains: the lack of objective methods to quantify the certainty of diagnostic assessments derived from these probability density functions (Begoli et al., 2019). Most validation workflows fail to utilize the morphological information within the posterior PDF density, masking the non-uniqueness of the solution (Motallebzadeh et al., 2017a, 2017b, 2018). To move toward defensible personalized medicine, we must move beyond predicting the state of a system and begin objectively quantifying the certainty of the prediction itself.

The clinical utility of patient-specific modeling depends on translating this complex uncertainty into actionable diagnosis. In a middle-ear context, consider a model-based diagnostic assessing whether a patient’s stapedial annular ligament stiffness is consistent with otosclerotic fixation. If the inferred posterior PDF spans both pathological and physiological ranges with comparable probability, no reliable surgical recommendation can be made. More broadly, the spread of a probability distribution directly represents the risk of diagnostic error; a wide or multimodal posterior PDF indicates that clinical data cannot uniquely resolve the patient’s physiological state (Abdar et al., 2021; Hüllermeier & Waegeman, 2021). Without an objective measure of certainty, model-based assessments remain opaque systems that may conceal critical parameter ambiguities, increasing the risk of misdiagnosis or unnecessary intervention (Ghassemi et al., 2020; Kompa et al., 2021). To be safely integrated into clinical workflows, these diagnostics must provide a clear confidence signal that identifies not only the most probable parameter state but also the certainty with which that state has been determined (Begoli et al., 2019).

SBI facilitates this personalized diagnosis by mapping clinical measurements to specific physiological parameters, offering higher interpretability than global physiological responses — such as middle-ear absorbance — where multiple pathologies may obscure one another (Tejero-Cantero et al., 2020). However, the reliability of SBI-derived posterior PDFs cannot be assumed from training performance alone. Hermans et al. (2022) demonstrated through extensive benchmarking that all major SBI algorithms — including neural posterior estimation variants — can produce overconfident posterior approximations that exclude plausible parameter regions, rendering them unreliable for scientific inference without further validation. A complementary and distinct concern is model misspecification: when the model fails to accurately represent the true physical system, the inferred parameter PDFs may be systematically wrong even when the inference algorithm performs (Schmitt et al., 2024). Beyond these algorithmic and structural failure modes, a third challenge arises from insufficient posterior PDF resolution: even a well-calibrated, correctly specified posterior PDF may remain too broad to support a clinical decision if measurement noise or limited training data prevents convergence to a diagnostically useful parameter range. Global calibration tools such as simulation-based calibration ([SBC]; Talts et al., 2020) address algorithmic validity averaged over the prior distribution, but cannot assess the resolution of inference for a specific patient observation — the very information required for clinical decision support.

These concerns are further compounded by “metric degeneracy,” wherein common scalar summaries fail to capture the underlying morphology of the posterior PDF (Abdar et al., 2021). In current practice, posterior PDF certainty is typically summarized by a single scalar — for example, the [2.5, 97.5]% credible interval width or standard deviation — which serves as the primary criterion for assessing parameter resolution and informing clinical decisions. However, such measures are inherently topology-blind: they assign identical scores to fundamentally different posterior PDF morphologies that carry very different diagnostic implications. A narrow, spike-like posterior and a broad plateau-like distribution with identical 95% mass width represent categorically different states of clinical certainty, yet are indistinguishable by this metric alone. This degeneracy is not limited to width-based measures; as established in our prior work on spectral biofidelity assessment, single-metric evaluations systematically mask structural features of complex frequency-response functions (FRFs) or spectral responses (Koshe et al., 2026). Moreover, in the parameter domain, a single width value cannot distinguish between a stable unimodal posterior — indicating unique resolution of a physiological state — and a bimodal distribution indicating that two distinct pathological states remain equally plausible (Hüllermeier & Waegeman, 2021). This blind spot is clinically consequential: a diagnostic system that cannot quantify certainty may report a confident parameter estimate even when the underlying posterior provides no meaningful resolution between normal and pathological states.

No existing framework provides a per-observation, multi-metric certainty signal for SBI-inferred parameter distributions that is simultaneously sensitive to posterior morphology, free of metric-specific blind spots, and applicable to clinical practice without additional forward simulations. To address this gap, we propose a **Composite Certainty Framework** for the multidimensional quantification of posterior PDFs in model-based machine learning that can extend to physics-informed applications. Building on our prior Composite Biofidelity Framework for spectral similarity assessment (Koshe et al., 2026), the present work extends the same rank-aggregation philosophy from the frequency-response functions to the inferred parameter posteriors. Five complementary uncertainty metrics — standard deviation, interquartile range, full width at half maximum, Shannon entropy, and 95% mass width — each sensitive to distinct features of posterior morphology, are aggregated using a non-parametric Borda count (Dwork et al., 2001) to produce a unit-free composite certainty score. This approach mirrors its recent adoption for robust consensus ranking in high-dimensional genomics, where no single feature importance measure is universally reliable across diverse data structures (Degenhardt et al., 2019). Critically, this framework operates entirely on the posterior samples already produced by the trained SBI estimator, requiring no additional forward simulations and no modification of the inference algorithm, making it directly deployable alongside existing SBI pipelines. We demonstrate the framework on the posterior distributions inferred by SBI for the parameters of a finite-element model of the human middle ear (Motallebzadeh et al., 2025), examining how composite certainty evolves as a function of measurement noise and training dataset size across seven physiological parameters. The results establish objective saturation thresholds for training dataset size requirements and noise tolerance limits, transforming SBI from a parameter estimation tool into a self-validating diagnostic system that reports its own confidence alongside its predictions — a capability essential for the responsible clinical deployment of model-based personalized medicine.

## II. METHODS

This section details the methodology for evaluating the certainty of inferred parameter distributions within an SBI-tuned finite-element (FE) middle-ear framework. Certainty is characterized by quantifying the convergence and dispersion of posterior probability distributions across two experimental dimensions: (1) the learning-curve behavior as training dataset size increases, which probes epistemic uncertainty arising from FEM simulation budget constraints; and (2) the noise-robustness behavior as Gaussian measurement noise is systematically added to FEM simulation outputs, which probes epistemic uncertainty arising from measurement artifacts (Motallebzadeh et al., 2025). Together, these two dimensions span the principal reducible uncertainty sources that govern whether an SBI-derived posterior achieves sufficient diagnostic resolution for clinical use.

### A. Reference Experimental Data

Reference measurements were obtained from a single cadaveric temporal bone (specimen TB24), serving as a controlled proof-of-concept for the proposed certainty framework. Data included wideband tympanometry for ear-canal input impedance and acoustic absorbance, alongside stapes velocity measurements for middle-ear transmission, providing a multivariate FRF target influenced by multiple physiological parameters simultaneously. Seventeen independent measurement sets were collected at ambient pressure under identical conditions to characterize the intrinsic test-retest variability of the preparation. The standard deviation of these repeated FRFs defines the experimental noise reference (σ) used to scale the noise levels in the robustness analysis (Section C), ensuring that the noise perturbations applied during evaluation are anchored to empirically observed measurement uncertainty rather than arbitrarily chosen values. These datasets serve as both the physiological fitting target for the SBI framework and the empirical reference for all certainty metric evaluations (Motallebzadeh et al., 2025).

### B. Finite-Element Model and SBI Framework

A deterministic finite-element model of the TB24 specimen was developed from high-resolution μCT images and implemented in COMSOL Multiphysics (Motallebzadeh et al., 2025). Seven free parameters, including cochlear load, structural damping, Young’s moduli (YM) of joints, ligaments, stapes annular ligament (SAL), tendons, and eardrum, were selected for inference due to their clinical relevance and influence on middle-ear dynamics; full details of the model geometry, prior specifications, and neural density estimator architecture are provided in Motallebzadeh et al. (2025).

The SBI framework (Tejero-Cantero et al., 2020) was trained on 10,000 pre-simulated FEM datasets, mapping randomly sampled prior parameter values to their corresponding acoustic and vibratory FRFs. The resulting amortized posterior estimator can be queried instantaneously for any observation without retraining. Once trained, the estimator was conditioned on the TB24 reference FRFs to generate 1,000 posterior samples per parameter for each simulated noise level and training dataset size evaluated in this study, forming the basis for all certainty metric computations in Section D.

### C. Experimental Design: Convergence and Noise Analyses

To evaluate the behavior of posterior certainty across the principal sources of reducible epistemic uncertainty, two computational sensitivity analyses were conducted, referred to hereafter as noise-robustness and convergence analyses. In both cases, the trained SBI estimator was queried on the TB24 reference FRFs and certainty metrics were computed from the resulting posterior PDFs.

In the **noise-robustness analysis**, the training dataset size was fixed at *N* = 10,000 simulations. Gaussian noise was added to the simulation outputs prior to SBI training at 21 logarithmically spaced multipliers ranging from 0.1σ to 10.0σ, where σ denotes the empirical standard deviation of the 17 repeated TB24 measurements defined in Section A. The lower bound (0.1σ) approximates near-ideal measurement conditions, while the upper bound (10.0σ) represents severe artifact levels well beyond realistic clinical acquisition, together spanning the full dynamic range of the framework’s noise tolerance.

In the **convergence analysis**, the noise level was held constant at σ and the SBI estimator was retrained at 21 logarithmically spaced training dataset sizes from *N* = 100 to *N* = 10,000 simulations. Logarithmic spacing was chosen because posterior PDF convergence is expected to follow a learning-curve trajectory that saturates progressively — linear spacing would over-sample the typically well-converged high-*N* regime and under-sample the critical low-*N* transition region where certainty changes most rapidly.

Both analyses were repeated across 10 independent random seeds related to the stochastic initialization of the SBI neural network, the random subsampling of training data at each dataset size, and the random noise realizations applied to simulation outputs. Certainty metrics were computed from 1,000 posterior samples per parameter for each noise level and training size evaluated. The two analyses are structurally symmetric — 21 conditions × 10 seeds each — making their composite certainty trajectories directly comparable.

### D. Certainty Metrics

To capture the multidimensional nature of posterior PDF dispersion and mitigate the metric degeneracy of singular scalar summaries, five complementary certainty metrics were selected, each sensitive to distinct distributional features. This multi-metric approach is motivated by the insufficiency of any single scalar summary for characterizing complex posterior morphologies as described in Introduction. All metrics were computed from *M* = 1,000 posterior samples {*θ*_ⱼ_}, *j* = 1, …, *M*, drawn from the SBI posterior for each of the seven inferred parameters.

#### 1. Spread and Variance Metrics

This class quantifies global dispersion of the posterior mass. ***Standard Deviation (STD)*** measures global dispersion around the sample mean *θ̅*:

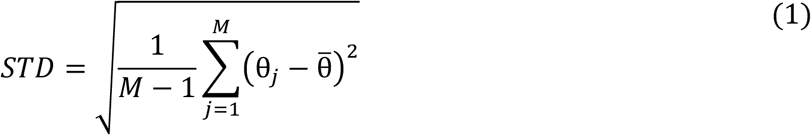

***Interquartile Range (IQR)*** measures the central 50% of the posterior mass, offering robustness to outliers and heavy tails:

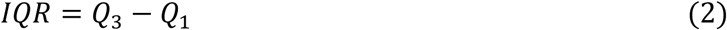

#### 2. Morphological and Resolution Metrics

For the following two metrics, the posterior density was estimated using kernel density estimation (KDE) with a Gaussian kernel and Scott’s bandwidth rule (Scott, 1992), evaluated on a uniform grid of K=1,000 points spanning the sample range. ***Full Width at Half Maximum (FWHM)*** is computed from the KDE of the posterior as the span between the outermost points where the estimated density exceeds half the global maximum. In multimodal posteriors this encompasses all modes above this threshold, reflecting the total high-probability span rather than primary peak sharpness alone:

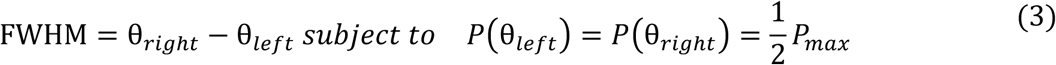

***Mass Width (95% Credible Interval)*** is the shortest interval containing 95% of the total posterior probability mass, providing a direct measure of diagnostic certainty bounds (Hyndman, 1996):

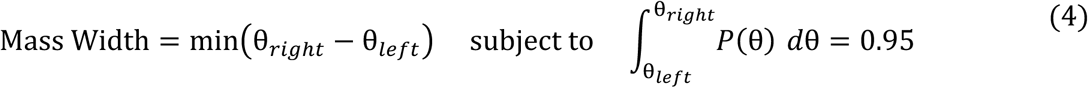

#### 3. Information-Theoretic Metric

***Shannon Entropy (****H)* quantifies the overall predictive uncertainty of the posterior. Unlike width-based measures, entropy is sensitive to the full morphology of the distribution, including multimodality and tail structure (Cover & Thomas, 2005). It is estimated via numerical integration of the KDE:

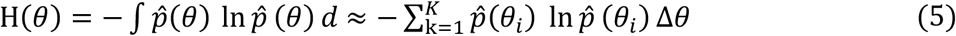

where *p^*(θ) is the KDE of the posterior samples, K = 1,000 is the number of grid points, and Δθ is the uniform grid spacing.

### E. Non-Parametric Rank Aggregation

To derive a single, unit-free consensus score that integrates information from the five complementary uncertainty metrics — each measuring posterior dispersion in its own physical unit — a non-parametric Borda rank aggregation procedure (Dwork et al., 2001) was applied **independently for each of the seven inferred parameters** and for each of the two experimental analyses (training-size convergence and noise robustness). Because Borda aggregation operates on ranks rather than raw values, the differing units and dynamic ranges of the underlying metrics are handled intrinsically, with no normalization of raw metric values required prior to aggregation. The procedure proceeds in three stages, with the final per-parameter outputs displayed in FIG 5.

#### 1. Stage 1 — Per-seed metric computation

For each of the 10 random seeds and each of the 21 experimental conditions (training sizes from 100 to 10,000, or noise levels from 0.1σ to 10σ), the 1,000 posterior samples for the parameter of interest were extracted and the five uncertainty metrics (IQR, STD, FWHM, Shannon entropy, mass width) were computed on those samples. This produces a 21 conditions × 5 metrics matrix of raw dispersion values per seed.

#### 2. Stage 2 — Within-metric ranking and rank summation

Within each metric column of the per-seed matrix, the 21 conditions were ranked from 1 (smallest dispersion → most certain condition for that metric) to 21 (largest dispersion → least certain condition for that metric). The five within-metric ranks per condition were then summed to produce a raw Borda score for that condition, with a theoretical range from 5 (ranked first by all five metrics, i.e., unanimous most certain) to 105 (ranked last by all five, i.e., unanimous least certain). Lower raw Borda scores thus indicate cross-metric consensus of greater certainty.

#### 3. Stage 3 — Cross-seed aggregation

For each condition, the raw Borda scores were aggregated across the 10 seeds, producing a final Borda curve summarized by the seed-mean with a 10th–90th percentile envelope quantifying seed-induced variability. Stages 1–3 were performed independently for each parameter and each experimental analysis, yielding 14 final Borda curves in total (7 parameters × 2 analyses), shown as the 7 rows × 2 columns layout in FIG. 5.

For visualization in FIG. 5, each per-parameter Borda curve (mean and percentile envelope jointly) was min-max scaled to a [0, 1] range, where **0.0 corresponds to the most certain and 1.0 to the least certain** within that curve. Across all three quantitative scales used in this framework — within-metric ranks (rank 1), raw Borda scores (5), and normalized display scores (0.0) — the most certain condition is consistently the one with the lowest value.

## III. RESULTS

### A. Evaluation of Metric Sensitivity and Degeneracy

#### 1. Divergence of Individual Certainty Metrics

To evaluate the discriminative power and inherent insensitivities of the five certainty metrics, twelve controlled PDF scenarios were applied against a reference Gaussian benchmark (FIG. 1). These scenarios were designed to represent pathological PDF morphologies that challenge scalar certainty summaries, including heavy-tailed distributions, multimodal topologies, and skewed or truncated masses. Each scenario was scaled so that the 95% mass width remained identical at 1.96 — the full width of the 95% credible interval of the reference Gaussian (σ = 0.5) — isolating each metric’s sensitivity to internal distributional morphology independent of overall spread. As shown in FIG. 1, fundamentally distinct PDF shapes yield identical credible interval widths, illustrating the metric degeneracy problem introduced in Introduction.

**FIG. 1.**
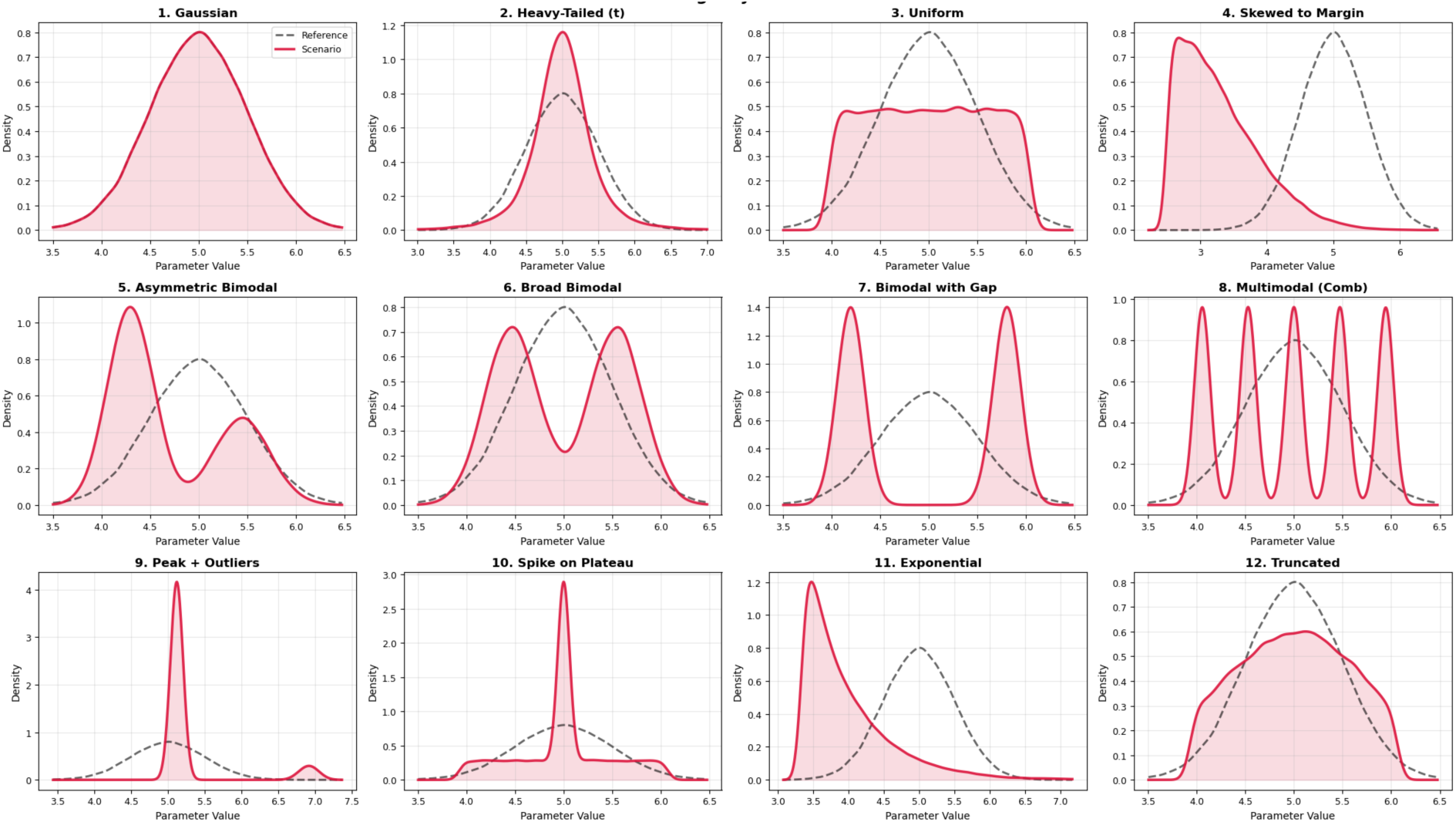
Twelve controlled PDF morphologies with identical 95% mass width (= 1.96, equal to the full width of the 95% credible interval of the reference Gaussian, σ = 0.5). Each scenario (red) is overlaid on the reference Gaussian (black dashed). Despite identical credible interval widths, the distributions span fundamentally different morphologies — from unimodal Gaussian to heavy-tailed, multimodal, and truncated shapes — demonstrating that mass width alone cannot distinguish between categorically different states of PDF certainty.

#### 2. Analysis of Metric Discrimination

To quantify how effectively each metric distinguishes among the twelve iso-width scenarios, we computed the coefficient of variation (CV) for each metric across the twelve morphology scenarios and visualized scenario-specific responses as a CV-weighted heatmap of min-max normalized scores (FIG. 2). Two complementary layers of information are encoded in this representation. First, the CV value reported in each column header quantifies the metric’s overall discriminative power across all twelve scenarios — a higher CV indicates that the metric assigns meaningfully different scores across morphologies, while a CV near zero indicates near-uniform response and thus limited global discrimination. Second, the color pattern within each column reveals the metric’s scenario-specific sensitivity profile: within a given column, scores are normalized so that 0.0 (yellow) corresponds to the scenario the metric identifies as most certain, and 1.0 (red) corresponds to the scenario it identifies as least certain, with 0.5 (orange) indicating no discrimination. Columns compressed toward orange throughout indicate a metric that cannot resolve differences between scenarios regardless of morphology.

**FIG. 2.**
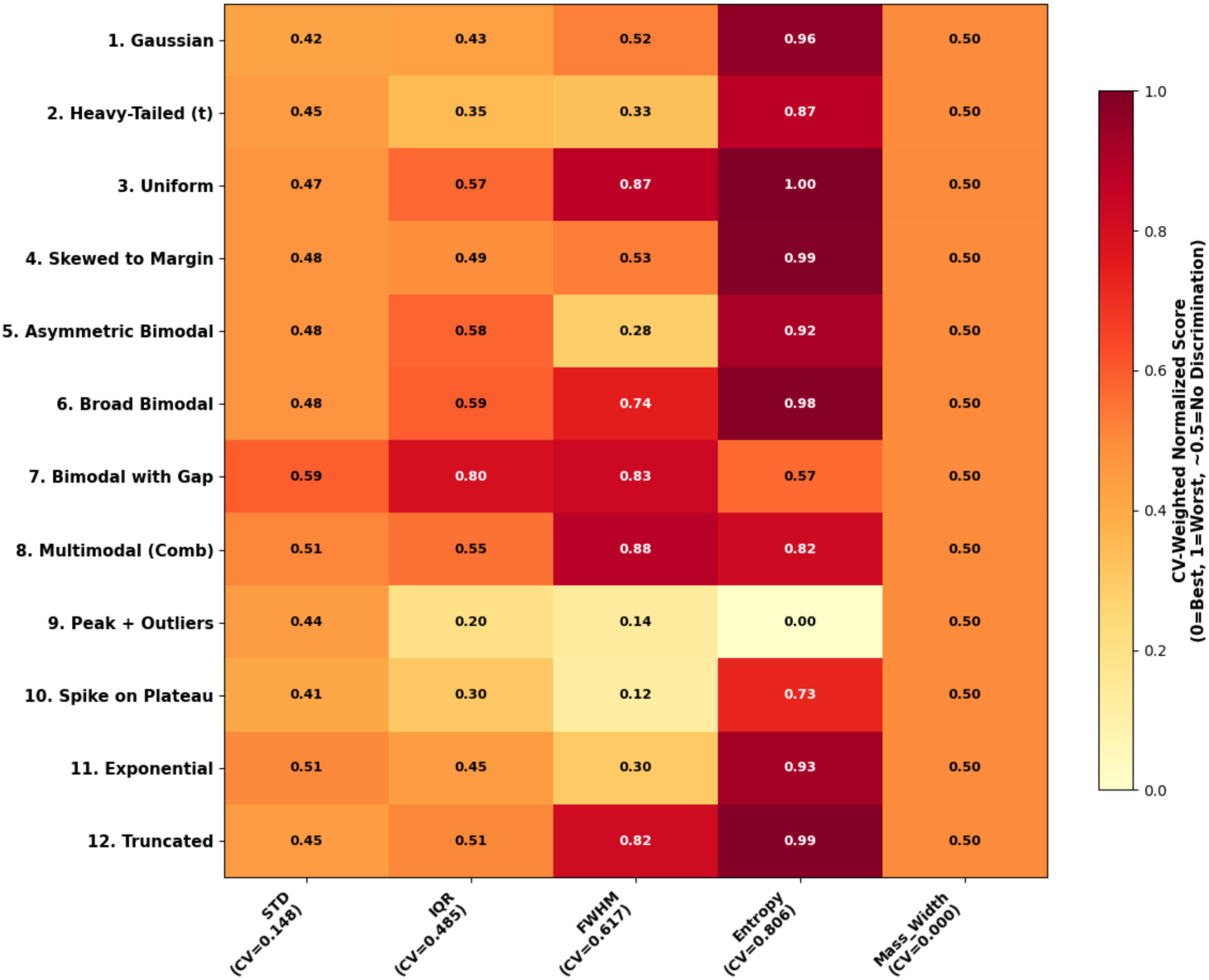
CV-weighted heatmap of normalized certainty metric responses across twelve iso-width PDF scenarios. Two layers of information are encoded: (1) the CV value in each column header quantifies overall discriminative power across all scenarios — higher CV indicates stronger global discrimination; (2) the color within each column reveals scenario-specific sensitivity, where 0.0 (yellow) = metric identifies most certain state, 1.0 (red) = least certain, and 0.5 (orange) = no discrimination. Columns compressed toward orange indicate metric blind spots. 95% mass width (CV = 0.000) produces a uniformly orange column by construction, directly illustrating metric degeneracy.

By construction, 95% mass width produced a perfectly uniform orange column (CV = 0.000), confirming its complete inability to distinguish among the iso-width cases and directly illustrating the metric degeneracy problem. STD showed similarly weak global discrimination (CV = 0.148), with all scenarios assigned near-identical scores. IQR showed moderate selectivity (CV = 0.495), responding strongly to bimodal distributions with large inter-modal gaps but remaining insensitive to spike-like and outlier-dominated morphologies. FWHM exhibited stronger global discrimination (CV = 0.617) and a distinct within-column pattern, strongly penalizing distributions that flatten or broaden the dominant peak while remaining insensitive to concentrated spike-on-plateau and peak-with-outlier cases. Shannon entropy showed the strongest overall discrimination (CV = 0.806), spanning the full normalized range from yellow to red and correctly identifying the uniform distribution as maximally uncertain and the spike-with-outliers case as most concentrated. Overall, no single metric discriminates strongly and consistently across all PDF morphologies — each exhibits characteristic blind spots determined by its sensitivity class — motivating the multi-metric aggregation framework of Section E.

### B. Evaluation of Parameter Posterior Resolution

FIG. 3 shows the evolution of posterior distributions and corresponding maximum *a posteriori* (MAP) estimates for the seven inferred parameters across noise levels from 0.1σ to 10.0σ (panel a) and training dataset sizes from N = 100 to N = 10,000 (panel b), each for a representative seed (consistent patterns were observed across all ten seeds). As noise increases, posterior PDFs progressively broaden and flatten, reflecting a growth in epistemic uncertainty and a transition away from high diagnostic resolution. The corresponding MAP estimates remain comparatively stable at low noise for most parameters but exhibit increasing drift, jitter, or abrupt shifts as noise grows, with the exact onset and pattern varying by parameter — indicating that point-estimate reliability degrades parameter-specifically rather than uniformly. Under training size variation (panel b), posterior PDFs narrow and sharpen with increasing N, reflecting convergence of the neural estimator from an uninformative prior toward a resolved parameter state. MAP estimates show larger fluctuations at low N for most parameters, consistent with the dependence of inference on the training size; these fluctuations generally diminish as N grows, though the rate of contraction varies by parameter.

**FIG. 3.**
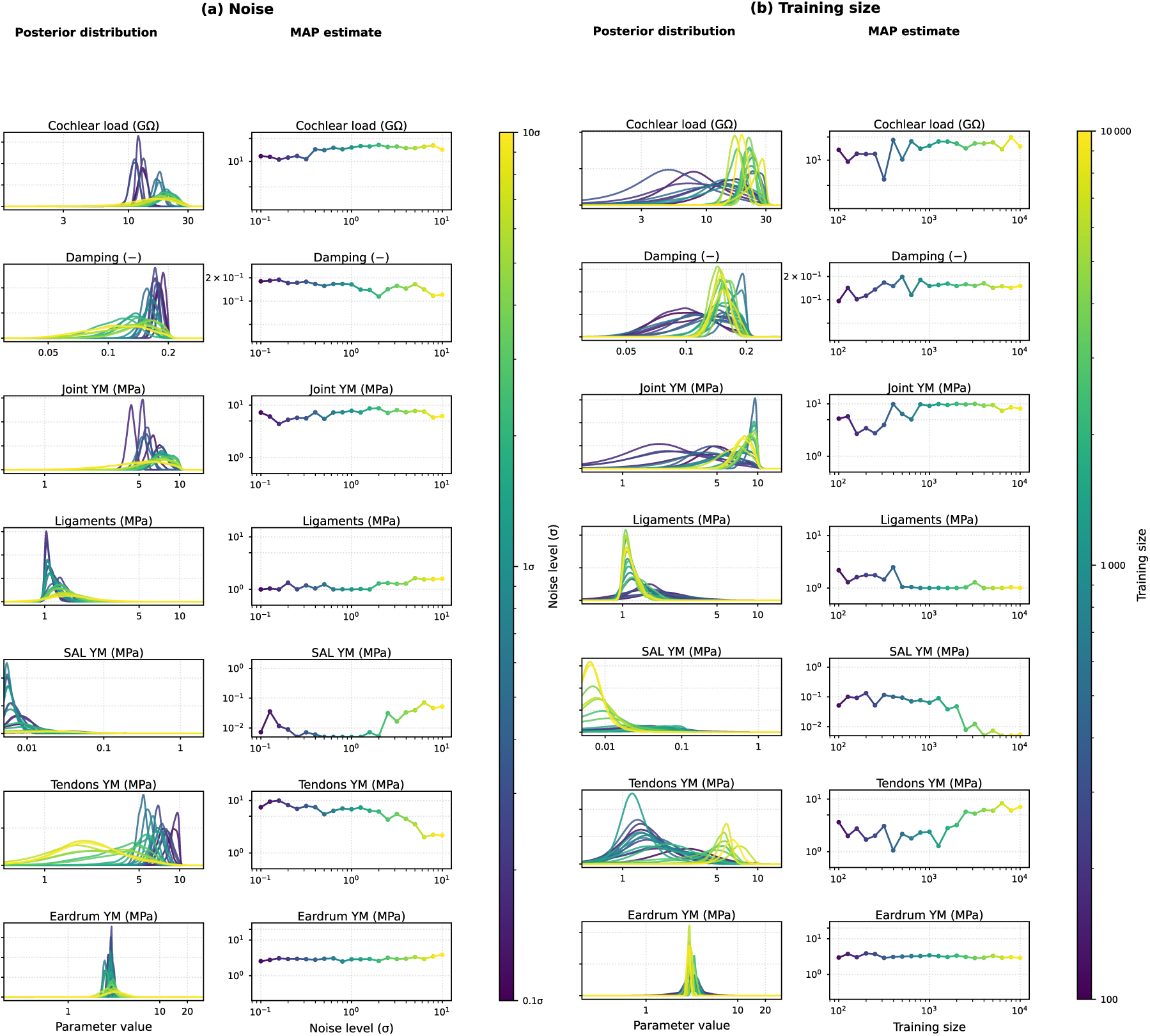
Parameter posterior evolution as a function of measurement noise (a) and training dataset size (b). In each panel, posterior PDFs (left columns) and MAP estimates (right columns) are shown for the seven inferred middle-ear parameters across 21 logarithmically spaced conditions. Panel (a): noise levels from 0.1σ to 10.0σ, where σ denotes the empirical standard deviation of the 17 repeated TB24 measurements (Section A); training size fixed at N = 10,000. Panel (b): training sizes from N = 100 to N = 10,000; noise fixed at σ. Colors encode the experimental condition via the colorbar. Results shown for one representative seed; similar patterns were observed across all ten seeds.

### C. Comparative Dynamics of Certainty Metrics

FIG. 4 presents the noise-response curves (panel a) and training convergence curves (panel b) for five certainty metrics applied to two contrasting parameters: SAL YM, which is the primary diagnostic target for otosclerosis and converges well under sufficient data, and Tendons YM, which is mechanically present but not linked to a specific pathology and exhibits persistently variable posterior PDFs (FIG. 3).

**FIG. 4.**
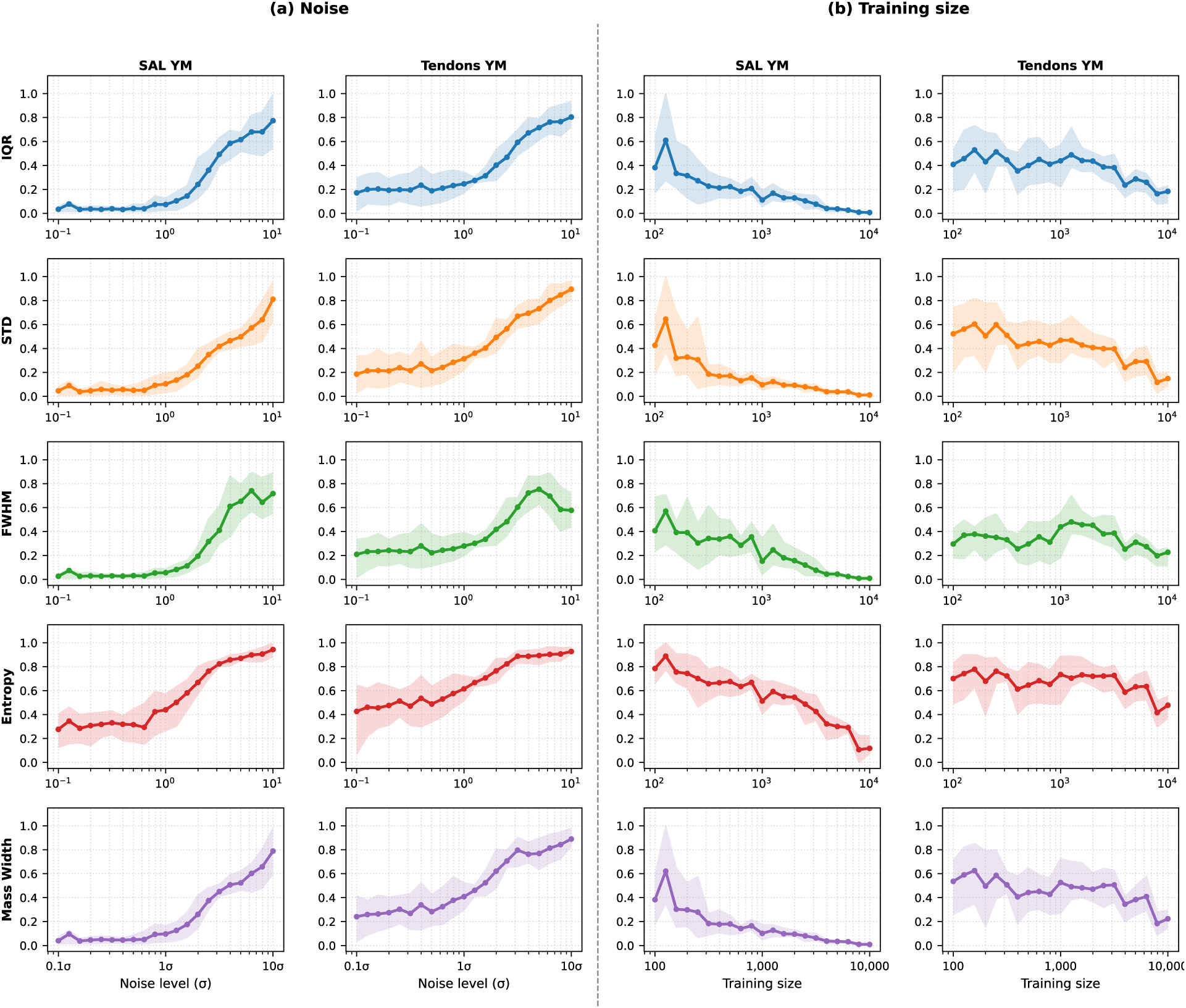
Certainty metric trajectories for SAL YM and Tendons YM as a function of measurement noise (a) and training dataset size (b). Each panel shows five certainty metrics (rows) for two contrasting parameters (columns): SAL YM, the primary diagnostic target for otosclerotic fixation, and Tendons YM, a mechanically present but pathologically non-specific parameter. All metric values are min-max normalized to [0, 1] across conditions per parameter, where 0 indicates minimum uncertainty (highest resolution) and 1 indicates maximum uncertainty. Solid lines denote the mean across 10 seeds and shaded bands indicate the 10th–90th percentile range. Panel (a): training size fixed at N = 10,000; noise varied from 0.1σ to 10σ. Panel (b): noise fixed at σ; training size varied from N = 100 to N = 10,000. Full trajectories for all seven parameters are provided in Supplementary Figures S1 (noise) and S2 (training).

In the noise analysis (panel a), SAL YM demonstrates the metric degeneracy identified in Section A in a clinically relevant context. Spread-based metrics — IQR, STD, FWHM, and mass width — remain near zero at low noise levels and rise steeply beyond approximately 1σ, consistent with a well-identified parameter whose posterior PDF remains concentrated until noise corrupts the physiological signal. Shannon entropy, however, starts at a substantially higher baseline at low noise and rises more gradually, indicating that even when the posterior PDF is narrow its internal morphology is not Gaussian — a feature invisible to spread metrics but captured by entropy. For Tendons YM, all metrics show a higher baseline and wider across-seed variability even at low noise levels, indicating that this parameter is poorly identified regardless of measurement quality, and FWHM exhibits a non-monotonic decline at high noise as the PDF peak structure collapses entirely.

In the training convergence analysis (panel b), SAL YM shows a sharp, consistent decline across all metrics from high uncertainty at N = 100 toward near-zero values at N = 10,000, confirming that this parameter is well-identified when sufficient training data are available and that all five metrics reach consensus on convergence. Tendons YM, by contrast, shows persistently elevated uncertainty with wide across-seed bands throughout the full training range, indicating that even the maximum available training size of 10,000 simulations does not produce a reliably resolved posterior for this parameter — a distinction with direct implications for the clinical utility of inference-based diagnostics.

The full per-metric, per-parameter trajectories are provided in Supplementary Figures S1 (noise) and S2 (training set size). Eardrum YM shows the highest posterior resolution, with all five metrics converging to near-zero under low noise and sufficient training. Cochlear Load, Joint YM, Ligaments YM, and Damping exhibit intermediate but broadly concordant metric responses. Notably, across all parameters, Shannon entropy maintains a monotonically increasing and well-behaved response even at very high noise levels where spread-based metrics become erratic. This reinforces the role of Shannon entropy as the most morphologically sensitive certainty indicator and further motivates the multi-metric aggregation approach.

### D. Composite Certainty Aggregation and Robustness Limits

To integrate the five individual metrics into a single consensus, the Borda rank aggregation procedure was applied to both dimensions we have explored. FIG. 5 presents the normalized composite certainty scores for all seven parameters as a function of measurement noise (panel a) and training dataset size (panel b).

**FIG. 5.**
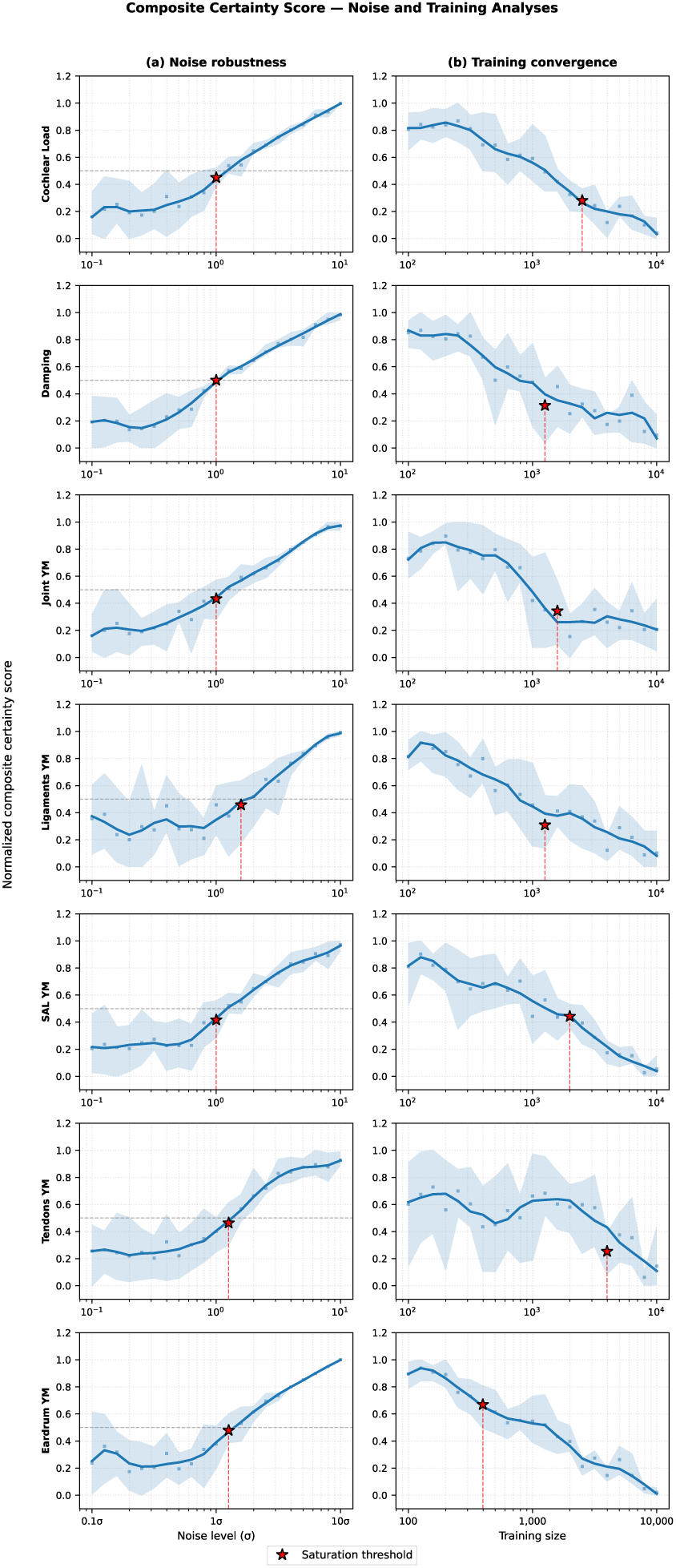
Composite certainty score as a function of measurement noise (a) and training dataset size (b) for the seven inferred middle-ear parameters. In each panel, the composite score is derived by Borda rank aggregation of five complementary certainty metrics across the ten random seeds. Scores are min-max normalized to [0, 1], where 0 indicates maximum posterior resolution and 1 indicates minimum resolution. Points show seed-averaged values, solid curves denote the smoothed consensus trajectory, and shaded bands indicate the 10th–90th percentile range across seeds. The red star marks the saturation threshold (score = 0.5), which identifies the maximum tolerable noise level (panel a) and the minimum sufficient training dataset size (panel b) for reliable parameter recovery. The dashed horizontal line at 0.5 indicates the saturation criterion.

In the noise analysis (panel a), all parameters exhibit a robustness plateau below approximately 1σ where the composite score remains near-zero, reflecting stable posterior resolution within the natural measurement variability. A non-zero baseline persists even at the lowest noise level, reflecting the irreducible aleatoric component of amortized inference. Beyond the plateau, a degradation slope emerges as the posterior PDF progressively broadens. The red star marks the saturation threshold — the noise level at which the composite score reaches 0.5, where 50% of diagnostic resolution is lost. This threshold falls at approximately 1σ for all seven parameters, establishing a consistent noise tolerance limit for the middle-ear model. This cut-off threshold could be adjusted based on the required certainty level in different applications.

In the training convergence analysis (panel b), the composite score begins near its maximum at N = 100 and decreases as the posterior PDF sharpens with increasing training data. The saturation threshold — here identifying the minimum sufficient training size — occurs between N = 1,259 and N = 1,585 for most parameters. Tendons YM is a notable exception, reaching the threshold only near N = 5,000, consistent with its limited identifiability discussed in Section C. The wider across-seed bands in panel (b) relative to panel (a) reflect stochastic dependence on the training data subset, confirming that training randomization is a meaningful source of uncertainty when determining required simulation budgets.

## IV. DISCUSSION

### A. Summary of Findings

This study established a multidimensional framework for quantifying posterior certainty in parameter inference, particularly in a SBI study. The benchmark analysis confirmed that standard scalar summaries are topology-blind: PDFs with identical credible interval widths differed radically in diagnostic implication, from a concentrated peak to a flat uninformative distribution. Shannon entropy and FWHM provided the strongest morphological discrimination, while STD — the most commonly reported uncertainty summary in biomechanical parameter estimation — showed the weakest sensitivity. Applied to the middle-ear model, the composite framework revealed parameter-specific identifiability profiles invisible to any single metric, with Shannon entropy maintaining robust monotonic sensitivity even where spread metrics became erratic — consistent with theoretical concerns about the limitations of width-based decompositions (Wimmer et al., 2023).

### B. Metric Degeneracy and Multi-Metric Consensus

The insufficiency of scalar uncertainty summaries has been recognized in machine learning (Hüllermeier & Waegeman, 2021), but its consequences for posterior parameter inference in biomechanical diagnosis have not been systematically characterized. Our results show this is not merely theoretical. For clinically critical parameters, entropy and spread metrics diverged substantially at low-to-moderate noise, meaning the diagnostic conclusion about parameter certainty depended entirely on the chosen metric. This behavior reflects the failure mode described by (Porthiyas et al., 2024), who showed that reporting the most likely parameters and their 95% credible interval can lead to confusing or misleading clinical interpretations. The Borda aggregation resolves this by requiring simultaneous consensus across complementary sensitivity classes: a parameter is deemed certain only when spread, morphological, and information-theoretic metrics agree, preventing any single blind spot from driving or biasing the diagnostic conclusion.

### C. Complementary to Existing SBI Diagnostics

The framework addresses a failure mode orthogonal to those targeted by existing SBI reliability tools. Hermans et al. (2022) showed that amortized SBI can produce overconfident posteriors that are too narrow. Our framework targets the complementary problem: posteriors that are valid in coverage but lack diagnostic resolution. Crucially, both can coexist — a posterior can pass global calibration checks (Talts et al., 2020) while still being uninformative for a specific clinical observation. The per-observation certainty score fills this gap. The non-zero uncertainty floor observed even at minimal noise reflects the irreducible aleatoric component of amortized inference. This is distinct from the reducible epistemic uncertainty captured by the training convergence analysis (Der Kiureghian & Ditlevsen, 2009; Kendall & Gal, 2017). A formal decomposition of these two sources within the composite score remains a direction for future work.

## V. CONCLUSION

We proposed a Composite Certainty Framework that addresses metric degeneracy in SBI-based biomechanical diagnostics. By aggregating five complementary metrics through non-parametric Borda rank voting, the framework produces a consensus certainty score robust to individual metric blind spots. It fills a gap that neither credible interval reporting nor global SBI calibration diagnostics can address: the per-observation question of whether a given posterior PDF provides sufficient resolution to support a clinical decision. The framework translates directly into actionable thresholds on measurement noise tolerance and minimum simulation budget, and is generalizable to any SBI-based personalized diagnostic pipeline where trustworthiness depends on posterior resolution rather than simply posterior existence. As Begoli et al. (2019) argued, principled uncertainty quantification is a prerequisite for clinical AI adoption — this framework provides that layer for physics-based inference, enabling a diagnostic system that knows when to withhold a conclusion rather than offer one that cannot be trusted.

## ACKNOWLEDGMENTS

This work was supported in part by the National Institutes of Health (NIH/NIDCD R21DC020274) and the Canadian Institutes of Health Research (PJT-189955). The authors would like to thank W. Robert J. Funnell for insightful discussions.

## AUTHOR DECLARATIONS

HM designed and conceived the study; HM and AK performed the simulations; HM, AK, and EST performed the inference and analyses; EST and KJ contributed to the conceptualization of the idea; HM wrote the manuscript; all authors contributed to the manuscript.

## DATA AVAILABILITY

The data and Finite Element model (in COMSOL ver 6.2) for this study are available from the corresponding author on request.

