## Supplementary Information for "Composite Certainty: Addressing Metric Degeneracy in Parameter Inference for Model-Based Diagnostics"

### Composite Certainty: Addressing Metric Degeneracy in Simulation-Based Biomechanical Diagnostics

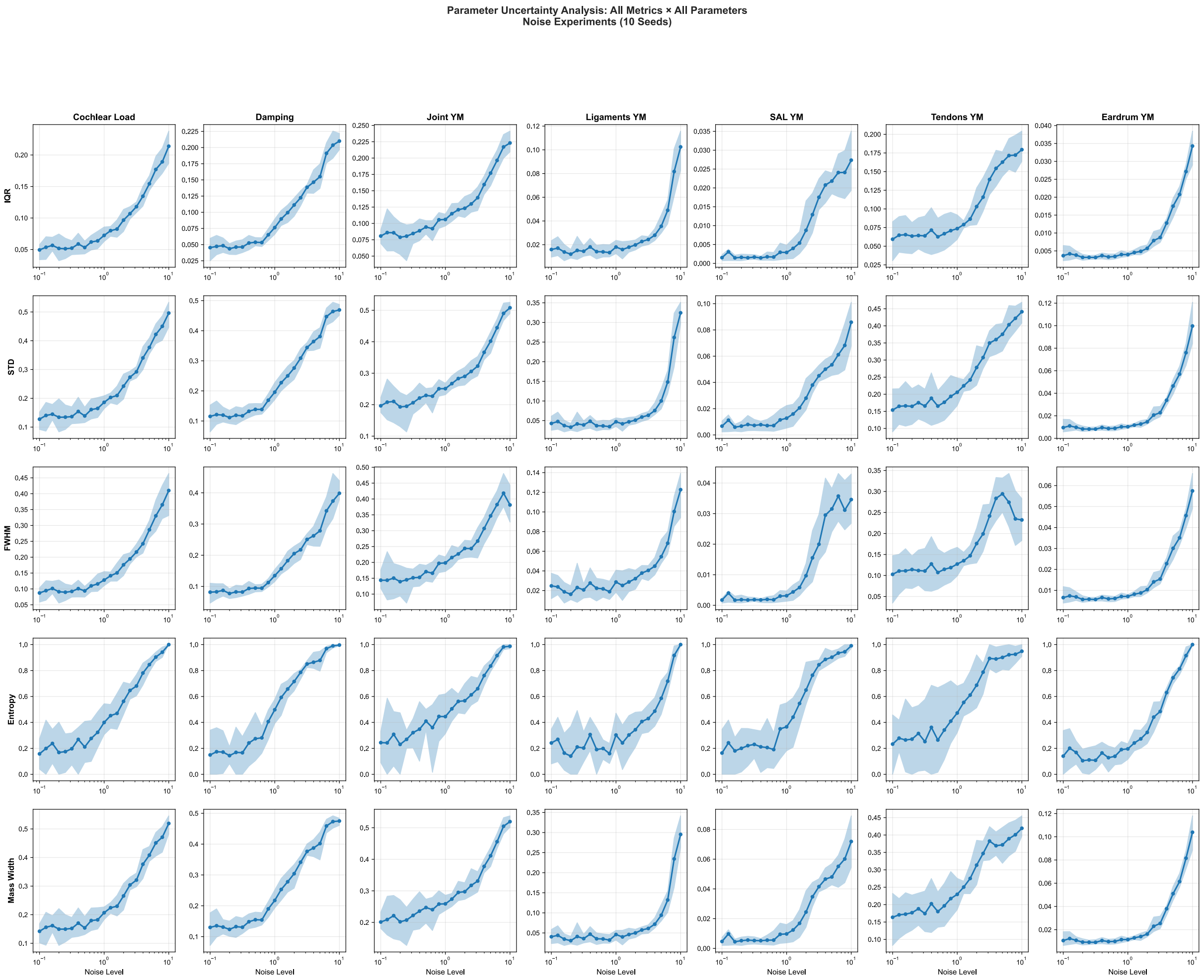


**Figure S1. Certainty metric trajectories for all seven inferred parameters as a function of measurement noise (0.1σ to 10σ).** Each panel shows the mean trajectory across 10 seeds (solid line) with the 10th–90th percentile band (shaded region). Rows correspond to the five certainty metrics (IQR, STD, FWHM, Entropy, Mass Width) and columns to the seven middle-ear parameters. Y-axis units reflect raw prior-normalized metric values and differ across parameters. SAL YM and Tendons YM are discussed in detail in the main text (FIG. 4); remaining parameters show intermediate but broadly concordant metric responses.

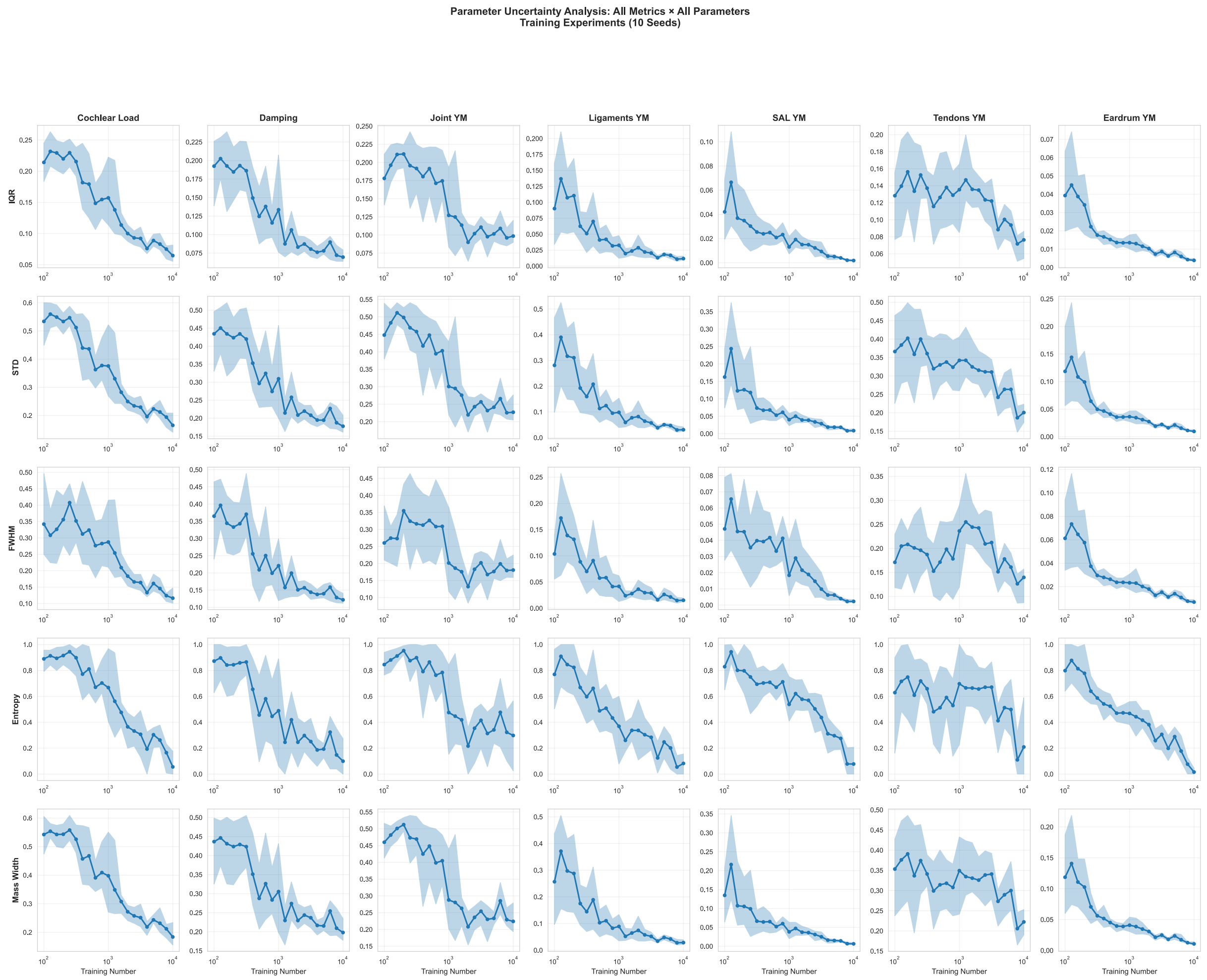


**Figure S2. Certainty metric trajectories for all seven inferred parameters as a function of training dataset size (N = 100 to N = 10,000).** Layout follows Supplementary Figure S1. Each panel shows the mean trajectory across 10 seeds (solid line) with the 10th–90th percentile band (shaded region). Wider across-seed bands relative to S1 reflect stochastic dependence on training data subsets rather than measurement noise. Eardrum YM shows the most consistent convergence across all five metrics; Tendons YM shows persistently elevated uncertainty throughout the full training range.
